# A small molecule stabilises the disordered native state of the Alzheimer’s Aβ peptide

**DOI:** 10.1101/2021.11.10.468059

**Authors:** Thomas Löhr, Kai Kohlhoff, Gabriella T. Heller, Carlo Camilloni, Michele Vendruscolo

**Affiliations:** Department of Chemistry, University of Cambridge, Cambridge, UK; Google Research, Mountain View, CA, USA; Department of Structural and Molecular Biology, University College London, London, UK; Dipartimento di Bioscienze, Università degli Studi di Milano, Milano, Italy

**Author notes:** Contributing authors.

**Keywords:** Markov model, molecular dynamics, disordered proteins

## Abstract

The stabilisation of native states of proteins is a powerful drug discovery strategy. It is still unclear, however, whether this approach can be applied to intrinsically disordered proteins. Here we report a small molecule that stabilises the native state of the Aβ42 peptide, an intrinsically disordered protein fragment associated with Alzheimer’s disease. We show that this stabilisation takes place by a dynamic binding mechanism, in which both the small molecule and the Aβ42 peptide remain disordered. This disordered binding mechanism involves enthalpically favourable local π-stacking interactions coupled with entropically advantageous global effects. These results indicate that small molecules can stabilise disordered proteins in their native states through transient non-specific interactions that provide enthalpic gain while simultaneously increasing the conformational entropy of the proteins.

## 1 Introduction

Drug development for Alzheimer’s disease has been a tremendous challenge in the past decades[1]. This condition is characterised by the formation of protein aggregates, such as fibrillar forms of the amyloid-β 42 peptide (Aβ42)[2, 3]. This protein fragment is intrinsically disordered, i.e. it does not form a single stable folded structure as a monomer, but instead exists in a dynamic equilibrium of states with transient local structure and fast transitions[4–12]. Many drug development efforts focused on aggregation-prone proteins such as Aβ42 attempt to target the already-formed fibril and/or the structurally elusive oligomeric species[13–15]. Other attempts aimed to identify small molecules capable of stabilising monomeric Aβ42 into a well-structured conformation[16–18] or generally interfering with the interaction of disordered proteins to structured partners by binding to their interfacing regions[19]. Since the most populated state of disordered proteins is conformationally highly heterogeneous, it has also been suggested that it may be more convenient to identify small molecules capable of stabilising this disordered state[20, 21]. The idea is that since the free energy landscape of disordered proteins is ‘inverted’ when compared with the funnel concept of folded proteins, with the disordered state as the free energy minimum and the ordered states exhibiting relatively high free energies [22], small molecules stabilising this minimum would be easier to develop, as they would not have to restructure the topology of the free energy landscape itself.

Independently from the strategy pursued, however, it is extremely challenging to characterize the binding mode of small molecules to disordered protein on an atomistic level. While some experimental methods such as nuclear magnetic resonance spectroscopy can provide quantitative information, it is often not sufficient to clearly understand the interactions and kinetics underlying the binding[20].

Molecular dynamics is one of the tools that can provide the necessary spatial and temporal resolution to study the interaction between disordered proteins and small molecules[20]. Together with Bayesian restraints from experimental data, molecular dynamics simulations have been used to characterize the thermodynamics of these binding modes in the case of the oncoprotein c-Myc[23] and Aβ42[5]. In the former study, urea was used as a control molecule to assess the sequence-specificity of the drug. In the latter case of Aβ42, we studied the interaction with the small molecule 10074-G5, and showed it was able to inhibit Aβ42 aggregation by binding the disordered monomeric form of the peptide. The interaction was characterized both experimentally, using various biophysical techniques, and computationally, using restrained molecular dynamics simulations with enhanced sampling. While in both systems the binding mode was found to be highly dynamic, a quantitative study of the kinetics was not possible due to the use of time-dependent restraints and biases during the simulation. The microscopic kinetics in form of contact lifetimes and autocorrelations could be especially instructive to fully understand the origin of entropic and enthalpic stabilisation in these extremely dynamic binding events (Fig. 1)[21].

**Fig. 1.**
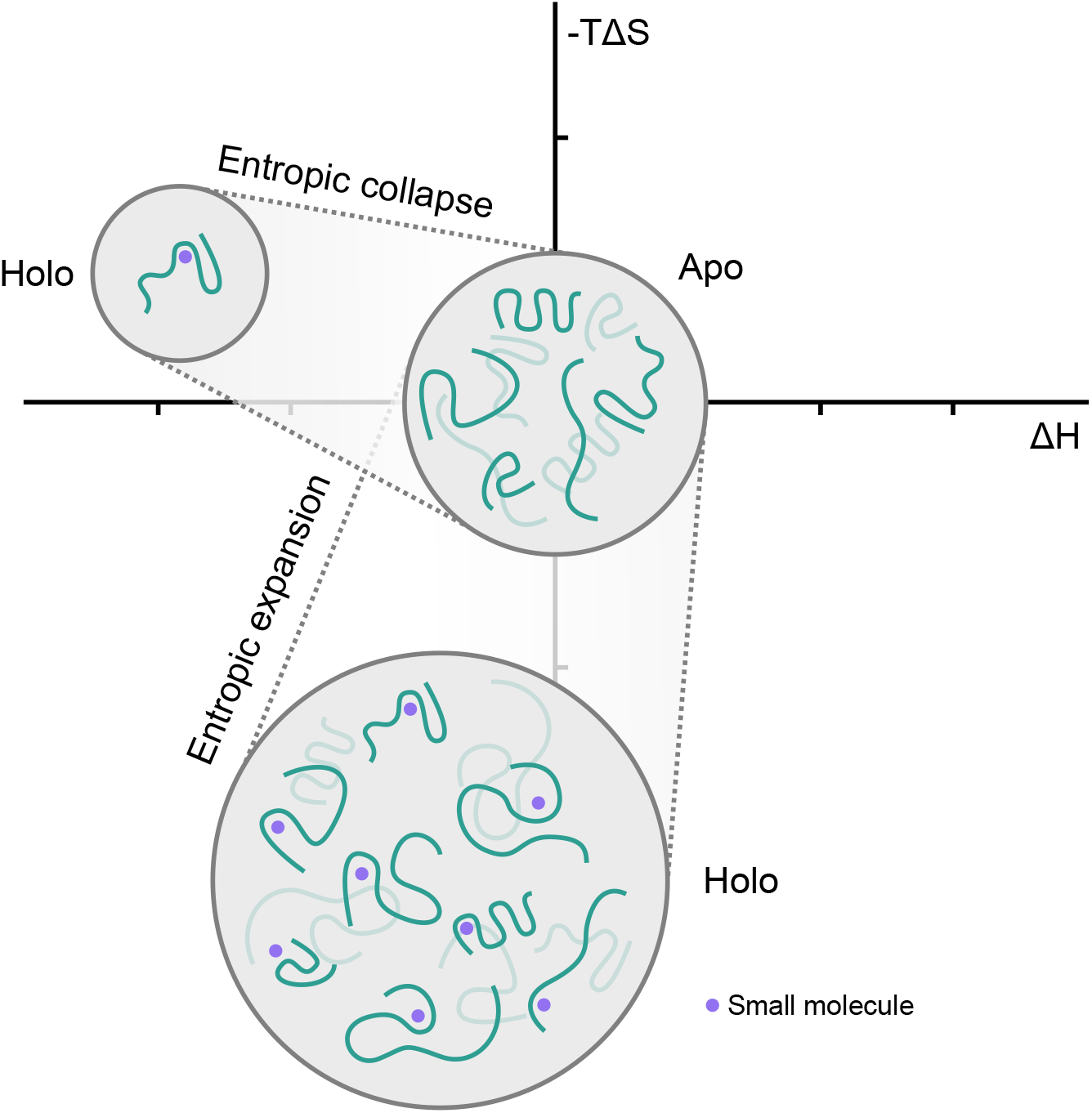
Illustration of two different native state stabilisation mechanisms of disordered proteins. The interaction with a small molecule can result in a reduction or increase of conformational space of the protein, thus resulting in a positive or negative entropic contribution to the binding free energy. A loss of entropic native state stabilisation will often be compensated with a stronger enthalpic binding affinity, while an increase in entropy often requires more dynamic and thus weaker binding.

A quantitative study of the kinetics of these interactions may allow a more targeted approach to the design of both drugs and better experiments to probe their binding modalities. However, even with atomistic computational approaches, gaining insight into the kinetics, i.e. transition rates, relaxation constants, autocorrelations, and state lifetimes can be challenging. This is because in contrast to folded systems, the definition of states for disordered proteins is not always clear: due to the generally shallow free energy landscape state transitions may be fast, but not always distinct[6]. New developments in the theory of dynamical systems now allow an optimal state decomposition and transition operator to be learned using deep neural networks, for example using the VAMPNet framework[24, 25]. To acquire kinetic information for a system one would traditionally use a Markov state model[26, 27]: One first finds a suitable low-dimensional embedding of the system coordinates, followed by using a clustering algorithm to define microstates. Transitions between these can then be counted to build up statistics and thus construct a transition matrix. This matrix can then be coarse-grained to obtain macroscopic kinetics[28, 29].

Koopman-operator[30, 31] based models such as VAMPNet combine these two steps into a single function that can be approximated by a neural network and also yield a probabilistic state assignment in lieu of a discrete one[24, 25]. The former feature has the advantage of both simplifying the model construction process, as the hyperparameter search over various dimensionality reduction and clustering techniques is replaced by a simplified search over neural network parameters, and allowing a more accurate model due to the use of a single arbitrarily non-linear function compared to two steps that are heavily restricted in terms of search space. Probabilistic state assignments are inherently well suited to disordered proteins, as the typically shallow free energy basins and low barriers can be encoded with some ambiguity. This constrained VAMPNet approach was recently utilized by us to determine the kinetic ensemble of the disordered Aβ42 monomer[6].

Here, we use this technique to build kinetic ensembles of Aβ42 with 10074-G5 and urea as a control molecule to expand on our previous thermodynamic ensembles[5]. We compare the transition rates, lifetimes, and state populations with the previous kinetic ensemble of the Aβ42 monomer[6], and further characterize the atomic-level protein-drug interactions.

## 2 Results

### 2.1 Molecular dynamics simulations and soft Markov state models

We performed two explicit-solvent molecular dynamics simulations of Aβ42 with one molecule of urea and one molecule of 10074-G5, respectively. Both simulations were performed in multiple rounds of 1,024 trajectories on the Google Compute Engine as described previously[6]. As before, we used a soft Markov state model approach using the constrained VAMPNet framework[24] to construct kinetic ensembles. The major advantages of this method, compared to regular discrete Markov state models, are the soft state definitions and the use of a single function mapping directly from arbitrary system coordinates to a state assignment probability, allowing for more optimal models. To aid our analysis, we added our previous simulation of Aβ42 with no additional molecules to our dataset. We refer to it as the *apo* ensemble[6]. We compared all ensembles using a decomposition into two states. In addition to being easier to interpret, this approach allows for a direct comparison of the slowest timescales in contrast to higher state-count models.

### 2.2 Computational and experimental validation

Constructing a kinetic ensemble using the constrained VAMPNet approach requires choosing the number of states and the model lag time. The latter is a critical parameter that needs to be chosen such that the model can accurately resolve both long and short timescales. This can be done by plotting the dependence of the slowest relaxation timescales on the lag time (Extended Fig. A1). A stricter measure is the Chapman-Kolmogorov test, comparing multiple applications of the Koopman operator estimated at a certain lag time *τ* with a Koopman operator estimated at a multiple of this lag time *nτ* (Extended Fig. A2, Methods)[32]. To evaluate sampling convergence, we visualized the dependence of the mean relaxation timescales on the number of trajectories used to evaluate these timescales (Extended Fig. A5). With sufficient sampling of kinetics, we would expect the global timescales to be unchanged within error. Experimental validation was performed by comparing back-calculated chemical shifts to ones from experiment[33]. Because the small molecule 10074-G5 only has minor effects on the chemical shifts of Aβ42[23], well below the error of the forward model, we used the chemical shift dataset from the apo ensemble as a point of comparison (Extended Fig. A3).

### 2.3 10074-G5 has minor impact on ensemble-averaged structural properties of Aβ42

To evaluate the influence of 10074-G5 and urea on the structural conformations of Aβ42, we calculated state-averaged contact maps and secondary structure content for each state of all ensembles (Extended Fig. A4a-c). In all cases we find a state decomposition into a more extended state with few inter-residue contacts, and a slightly more compact form with a higher number of local backbone interactions. We will refer to these as the compact and extended states, respectively. The addition of a small molecule has little effect on the formation of contacts and other structural motifs. This finding is consistent with our recent experimental thermodynamic and kinetic characterization of this interaction, and the absence of strong chemical shift perturbations in the holo ensemble[5].

### 2.4 10074-G5 and urea decelerate the formation of more compact states

Compared to the previously published kinetic ensemble of the apo form of Aβ42, the kinetic ensembles in the presence of both urea and 10074-G5 show a deceleration of more compact state formation (Fig. 2). Notably, the transition from the more compact form to the more extended state is unaffected. This change is also mirrored in the state populations, which exhibit a strong shift towards the extended state. We note that even though there are strong changes in the state populations, the ensemble-averaged contact maps are very similar (Extended Fig. A4a-c). This is likely due to the high sensitivity of the VAMPNet method to minor changes in free energy barrier regions. These will have a significant effect on the kinetics and thus state populations, but not on the ensemble averaged structure due to the relatively low thermodynamic weight[34]. While the lifetimes of the extended states increase, the ones for the more compact form are unchanged within model error. We can thus conclude that within our model, the small molecule has a strong effect on the contact formation rates, but no influence on the contact dissociation rates.

**Fig. 2.**
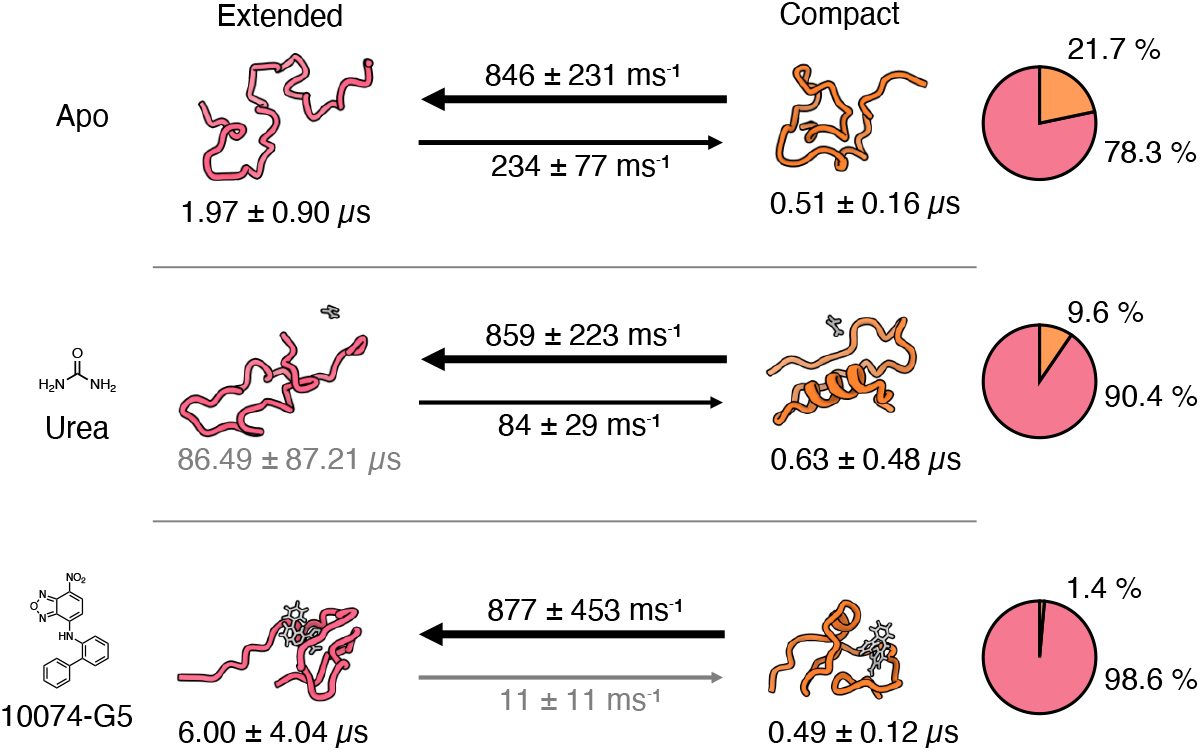
Impact of small molecules on the state transition rates, state lifetimes, and populations. The arrows indicate the mean state transition rates, the number below the representative structures is the mean state lifetime, the pie charts show the mean state populations. Errors are the standard deviations of the bootstrap sample of the mean over all 20 models.

### 2.5 Small molecules shift the system to more entropically stable states by short-lived local interactions

To evaluate the impact of 10074-G5 on the conformational space of Aβ42, we calculated the Ramachandran and state entropy for all ensembles, as well as the autocorrelation of sidechain χ_1_ dihedral angles (Fig. 3). The Ramachandran entropy can indicate relative flexibility of the backbone, thus revealing potential regions of dynamic changes as a result of interactions between the peptide and small molecule[5]. Resolving this change in the entropy over residues (Fig. 3a) indicates strong increases in the relatively hydrophobic C-terminal region of Aβ42. This entropy increase is confirmed globally by the sum of the entropies over all residues (Fig. 3b). As an alternative metric, we also calculated the entropy in the state assignments (Fig. 3c), this can be thought of as indicating the overall ambiguity in the state definition. Again, we find a relatively strong increase in the conformational entropy of Aβ42 for the ensemble with 10074-G5, and only minor increases for urea. These results are in agreement with our previous observations from simulations of the equilibrium ensembles in that the presence of 10074-G5 increases the conformations available to Aβ42, via the entropic expansion mechanism[5, 21].

**Fig. 3.**
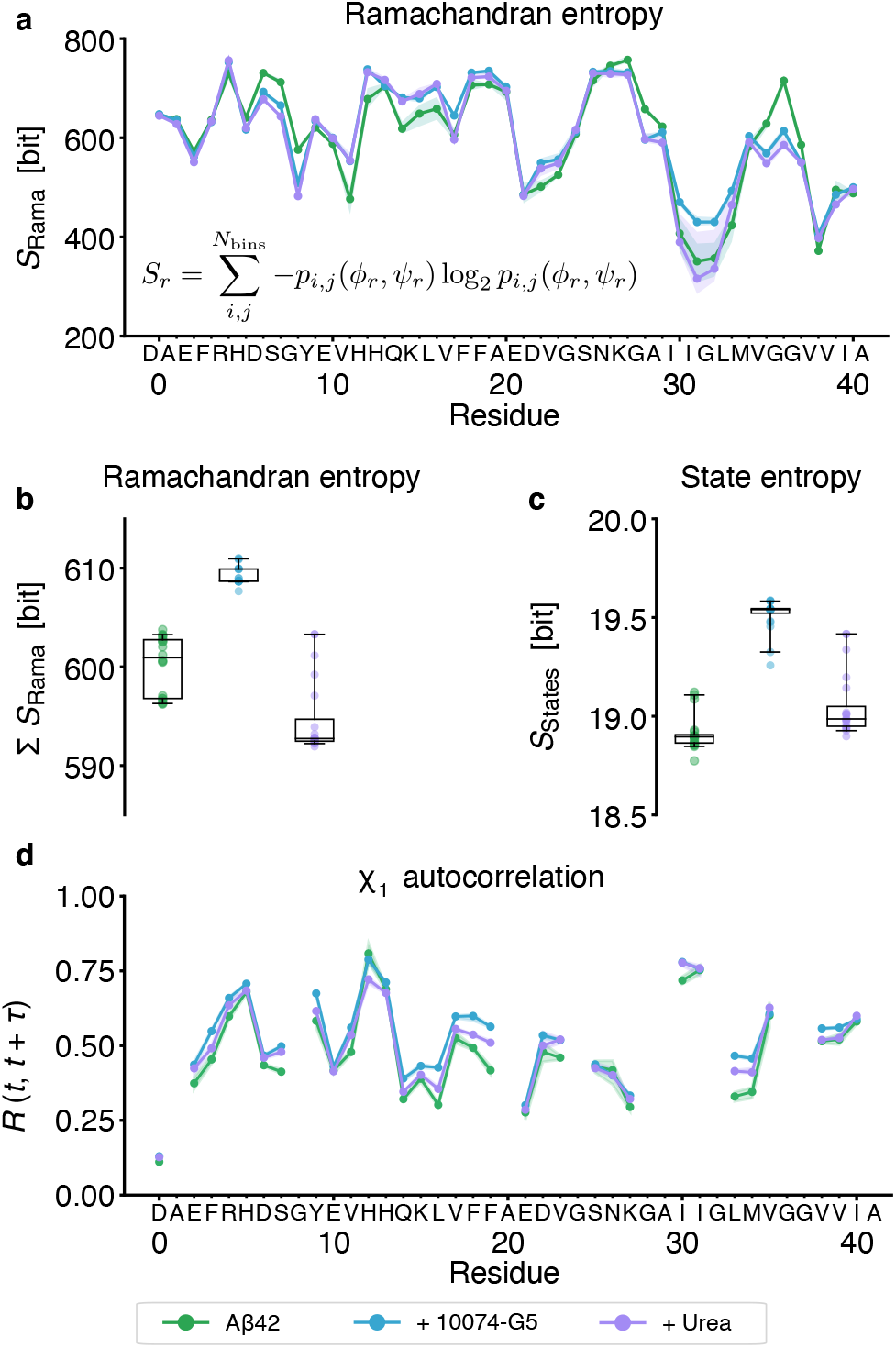
Effect of small molecules on conformational and state entropy of Aβ42, showing that 10074-G5 increases the conformational entropy of the peptide. **a** Ramachandran entropy, i.e. information entropy over the distribution of φ and ψ backbone dihedral angle conformations, using 100 bins. **b** Sum of the Ramachandran entropies over all residues for all ensembles. **c** State entropy, i.e., the population-weighted mean of the information entropy of each set of state assignments. More ambiguity in the state assignments leads to a correspondingly higher state entropy. **d** Autocorrelation of all sidechain χ_1_ dihedral angles with a lag time of *τ* = 5 ns. Shaded areas in **a** and **d** indicate the 95th percentiles of the bootstrap sample of the mean over all 20 models. Whiskers and boxes in **b** and **c** indicate the 95th percentiles and quartiles of the bootstrap sample of the mean over all 20 models, respectively.

To better understand the impact of the small molecule on local kinetics we calculated the autocorrelation of the sidechain χ_1_ dihedral angles (Fig. 3). We see an increase in the autocorrelation, specifically for aromatic residues and M35, indicating a slowing of side chain rotations. This suggests that despite an increase in the backbone entropy, the peptide is able to visit many locally stable states, resulting in local enthalpic stabilisation.

### 2.6 Interactions of 10074-G5 with Aβ42 are dominated by π-stacking and other electrostatic effects

To better understand the origin of the global and local effects of 10074-G5 on the ensemble we analysed the interactions on a residue and atomistic level (Fig. 4). While the probability of forming a contact between the small molecule and a residue shows certain mild preferences (Fig. 4a), these become more evident when looking at the lifetimes of these contacts (Fig. 4b). Here, the longest contacts are formed by π-stacking with certain aromatic residues (F4, Y10, F19, F20) and by interactions with M35. This result also explains the reduction in side-chain rotations for these residues (Fig. 3d). On an atomistic level the π-π interactions exhibit some anisotropy (Extended Fig. A6). The importance of the nitro- and benzofurazan fragments is also highlighted. Finally, we also investigated the conditionality of π-π interactions, i.e., if we see an interaction between the molecule and residue *i*, what is the probability of also observing an interaction with residue *j* (Fig. 4e-g)? The probabilities here are uniformly low but indicate a slight preference (13 %) for a triple π-stack involving the terminal aromatic ring of 10074-G5 and residues F19 and F20 of Aβ42. The importance of π-π stacking interactions was also noted in a computational study on the interactions of small molecules with α-synuclein[35].

**Fig. 4.**
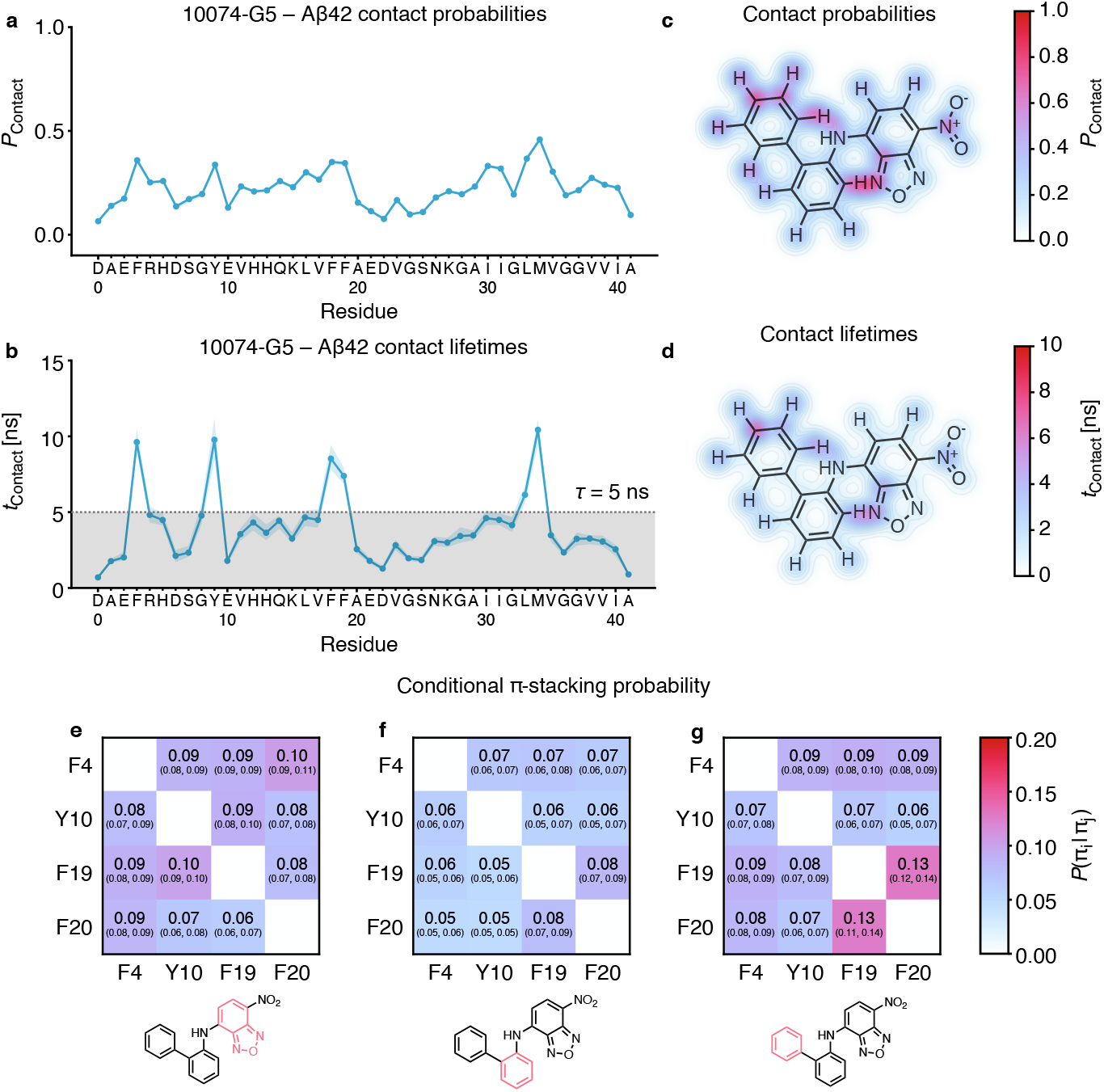
Residue- and atomic level interactions of 10074-G5 with Aβ42 showing regions on the small molecule responsible for binding. **a** Contact probabilities of 10074-G5 and Aβ42 with a cut-off of 0.45 nm. **b** Lifetimes of these contacts, estimated using a Markov state model for each contact formation with a lag time of *τ* = 5 ns, indicated with grey shading. Coloured shaded areas in **a** and **b** indicate the 95th percentiles of the bootstrap sample of the mean over all 20 models. **c**, **d** Contact probabilities and lifetimes of each atom of 10074-G5 with any residue of Aβ42. **e**, **f**, **g** Conditional probability of forming a π-stacking interaction, given an existing π-stacking interaction for all aromatic groups in the small molecule. Tuples indicate the 95th percentiles of the bootstrap sample of the mean over all 20 models.

These results indicate that this disordered binding mechanism operates on two levels whereby local enthalpically favourable interactions coupled with global entropically advantageous effects. The local interactions are predominantly of electrostatic nature and result in a reduction of sidechain rotations on specific residues. At the same time, these interactions also allow the exploration of more backbone conformations, thus resulting in a net entropy increase for Aβ42. This influence expands into the global kinetics of the system, significantly slowing the formation of local structure.

## 3 Discussion

The results outlined above present a possible example of the previously proposed entropic expansion mechanism for the binding of small molecules to disordered proteins[21, 36]. This mechanism is distinct from the entropic collapse and folding-on-binding mechanisms[37, 38]. The concept of disordered binding is difficult to probe, as the tools suitable to detecting small changes in the conformational ensemble of disordered proteins are still in their infancy[20]. Nuclear magnetic resonance experiments can provide information, but it should usually be interpreted in a structural framework, necessitating molecular simulations with ensemble-averaged restraints[39], or re-weighting approaches[40]. This constraint causes issues whenever we are also interested in kinetics, as by enhancing the sampling we modify the natural dynamics of the system. Nevertheless, an approach to incorporate ensemble-averaged experimental measurements into Koopman models has recently been proposed[41]. Neither is it generally possible to use enhanced sampling methods to study kinetics without having some *a priori* knowledge of the system states. A framework allowing the incorporation of experimental data into a kinetic model and also allowing the use of enhanced sampling methods such as metadynamics[42], without prior knowledge of states, would make the study of these systems easier and more accurate.

As we have shown, a kinetic model is crucial to fully explain the nature of these binding interactions. This is in part due to the ability to use the slowest timescales of the system to reliably define metastable states, something that is notoriously difficult for disordered proteins without access to the time dimension. This clustering alone is already sensitive enough to reveal differences between systems that are nearly invisible when comparing ensemble-averaged results and more conventional clustering methods[5]. Increases in local autocorrelation and global state transitions might be seen as indicators of both local enthalpic stabilisation and global entropic expansion. The former result hints at the possibility of designing small molecules that exhibit high specificity, as the global entropic stabilisation effect may be due to transient, local, enthalpically-favourable interactions[21]. The two level global entropy – local enthalpy effect becomes especially visible when looking at the timescales: The slowest state transitions of the protein are on the order of microseconds, while the local, enthalpically-favourable π-π interaction lifetimes are no longer than tens of nanoseconds.

The observed binding mechanism also identify π-π stacking interactions as a major driving force. Similar effects have been observed for the binding of another small molecule, fasudil, and α-synuclein, which is also intrinsically disordered[35]. We note that while that study proposed a ‘shuttling model’ mechanism to explain the diffusion of the small molecule on the α-synuclein surface, here we demonstrate the stabilisation of a native state of a disordered protein by a disordered binding mechanism. The π-π stacking phenomenon also plays a major role in liquid-liquid phase separation[43], suggesting a possible link between the effect of these small molecules and the hypothesized state of some proteins in a crowded environment. For molecular simulations, the force field may present a barrier in studying π-π interactions in more detail. This is because these interactions are not explicitly part of the potential, but only approximated with a combination of electrostatic and hydrophobic terms[44]. Polarizable force fields may offer a computationally affordable alternative that could more accurately model this type of binding[45].

Looking forward, it may become possible to pursue a drug discovery strategy for disordered proteins based on the stabilisation of their native states through the disordered binding mechanism that we have described here. This strategy would extend an approach to disordered proteins that has already proven successful for folded proteins[46], and would have the advantage of maintaining the proteins in their native functional states.

## 4 Methods

### 4.1 Details of the simulations

All simulations were performed on the Google Compute Engine with n1-highcpu-8 preemptible virtual machine instances, equipped with eight Intel Haswell CPU cores and 7.2 GB of RAM. Molecular dynamics simulations were performed with GROMACS 2018.1[47], with 1,024 starting structures sampled from the previously performed apo simulation[6] using the Koopman model weights. Each conformation was placed in the center of a rhombic dodecahedron box with a volume of 358 nm^3^, and the corresponding small molecule was placed in the corner of the box. The force field parameters for urea were taken from the CHARMM22* force field[48] and the ones for 10074-G5 were computed using the Force Field Toolkit (FFTK)[49] and Gaussian 09[50], as described previously[5]. The systems were then solvated using between 11,698 (11,707) and 11,740 (11,749) water molecules. Both systems were minimized using the steepest descent method to a maximum target force of 1,000 kJ/− mol/nm. Both systems were subsequently equilibrated, first over a time range of 500 ps in the NVT ensemble using the Bussi thermostat[51] and then over another 500 ps in the NPT ensemble using Berendsen pressure coupling[52]. In both equilibrations position restraints were placed on all heavy atoms. All production simulations were performed using 2 fs time steps in the NVT ensemble using the CHARMM22*[48] force field and TIP3P water model[53] at 278 K and LINCS constraints[54] on all bonds. Electrostatic interactions were modelled using the Particle-Mesh-Ewald approach[55] with a short-range cutoff of 1.2 nm. All simulations used periodic boundary conditions. We again used the fluctuation-amplification of specific traits (FAST) approach[56] to adaptive sampling, with clustering performed through time-lagged independent component analysis (TICA)[57, 58] using a lag time of 5 ns and Cα distances fed to the *k*-means clustering algorithm[59] to yield 128 clusters. 1,024 new structures were then sampled from these clusters based on maximizing the deviation to the mean Cα distance matrix for each cluster and maximizing the sampling of the existing clusters, using a balance parameter of *α* = 1.0, with all amino acids weighted equally. This approach was performed once for each ensemble, however we also chose to perform 32 additional long-trajectory simulations for the 10074-G5 ensemble, yielding a total of 2,079 trajectories for the latter, and 2,048 trajectories for the urea ensemble. The total simulated times were 306 μs and 279 μs for the 10074-G5 and urea ensembles, respectively. The shortest and longest trajectories for 10074-G5 (urea) were 21 (24) ns and 1,134 (196) ns. All trajectories were subsampled to 250 ps timesteps for further analysis.

### 4.2 Details of the neural networks

State decomposition and kinetic model construction was performed using the constrained VAMPNet approach[24, 25], using the same method described previously[6]. We again chose flattened inter-residue nearest-neighbour heavy-atom distance matrices as inputs, resulting in 780 input dimensions. We used the self-normalizing neural network architecture[60] with scaled-exponential linear units, normal LeCun weight initialization[61] and alpha dropout. We chose an output dimension of 2, thus yielding a soft two-state assignment. The datasets were prepared by first creating a test dataset by randomly sampling 10 % of the frames. In the case of 10074-G5 we excluded all frames in which the closest distance between the small molecule and peptide was higher than 0.5 nm. We then created 20 randomized 9:1 train-validation splits to allow a model error estimate. Training was performed by using three trials for each train-validation split and picking the best performing model based on the VAMP2 score[31] of the test set. We implemented the model using Keras 2.2.4[62] with the Tensorflow 2.1.0[63] backend. We chose the following model hyperparameters based on two successive coarser and finer grid searches: A network lag time of 5 ns, a layer width of 512 nodes, a depth of 2 layers, an L2 regularization strength of 10^−7^ and a dropout of 0.05. Training was performed in 10,000 frame pairs using the Adam minimizer[64] with a learning rate of 0.05, *β*_2_ = 0.99 and epsilon of 10^−4^, and an early stopping criterion of a minimum validation score improvement of 10^−3^ over the last five epochs. For the constrained part of the model, we reduced the learning rate by a factor of 0.02. We used a single Google Compute Engine instance with 12 Intel Haswell cores, 78 GB of RAM, and an NVidia Tesla V100 GPU.

### 4.3 Details of the kinetic analysis

After training, VAMPNet yields a state assignment vector *χ*(**x**_*t*_) for each frame **x**_*t*_ of the ensemble. Based on this vector, we can calculate state averages *A_i_* for any observable *A*(**x**_*t*_):

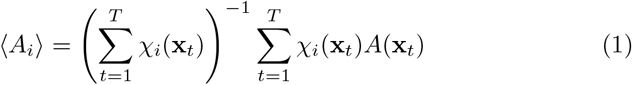

Here, *i* is the corresponding state and the sum runs over all time steps. To calculate an ensemble average ⟨*A*⟩, one first calculates a weight *w_t_* for each frame using the model equilibrium distribution *π*:

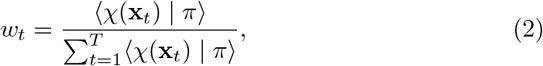

which leads to the ensemble average

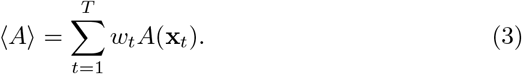

Because each trained model will classify the states in an arbitrary order, we need to sort the state assignment vectors based on state similarity. We did this by comparing the state-averaged contact maps using root-mean-square deviation as a metric, and grouping states based on the lowest value. Any deviations are thus accounted for in the overall model error.

### 4.4 Model validation

The Koopman matrix **K**(*τ*) is given directly by the neural network model, along with the equilibrium distribution *π*. We validated our models using the Chapman-Kolmogorov test:

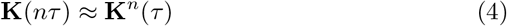

where *τ* is the model lag time, and *nτ* is a low integer-multiple of the lag time. The model should therefore behave the same way whether we estimate it at a longer lag time or repeatedly apply the transfer operator. We first estimate a suitable lag time *τ* by plotting the relaxation timescales over the chosen lag time. The lag time *τ* should be chosen to be as small as possible, but large enough to not have any impact on the longer relaxation timescales, which represent the slowest motions of the system. The temporal resolution of the model is thus given by this lag time. The relaxation timescales *t_i_* are calculated from the eigenvalues *λ_i_* of the Koopman matrix **K**(*τ*) as follows:

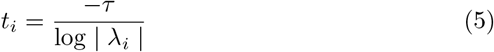

We can similarly compute the state lifetimes 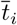 from the diagonal elements of the Koopman matrix **K**(*τ*)_*ii*_ using:

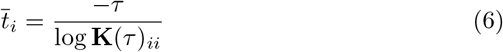

### 4.5 Experimental validation

We backcalculated the nuclear magnetic resonance chemical shifts using the CamShift algorithm[33] as implemented in PLUMED 2.4.1[65, 66]. We again used the same ensemble averaging procedure described above.

### 4.6 Errors

Errors are calculated over all trained neural network models. To obtain a more meaningful estimate, we only consider frames that were part of the bootstrap training sample of the corresponding model, i.e., one of the 20 models described above. The reported averages are the mean, and the errors the 95th percentiles over all 20 models, unless reported otherwise.

## Acknowledgements

G.T.H is supported by the the Rosalind Franklin Research Fellowship at Newnham College, Cambridge and the Schmidt Science Fellowship. We would like to thank Google and the Google Accelerated Science team for providing access to the Google Cloud Platform for simulations and analysis.

## Declarations

### 4.7 Code availability

Analysis code and notebooks are available from https://github.com/vendruscolo-lab/ab42-g5-ensemble and https://zenodo.org/record/5659241.

### 4.8 Data availability

Subsampled trajectory and intermediate data, as well as the trained neural network weights and analysis notebooks, are available from https://zenodo.org/record/5659241.

## APPENDIX A Extended Data

**Fig. A1.**
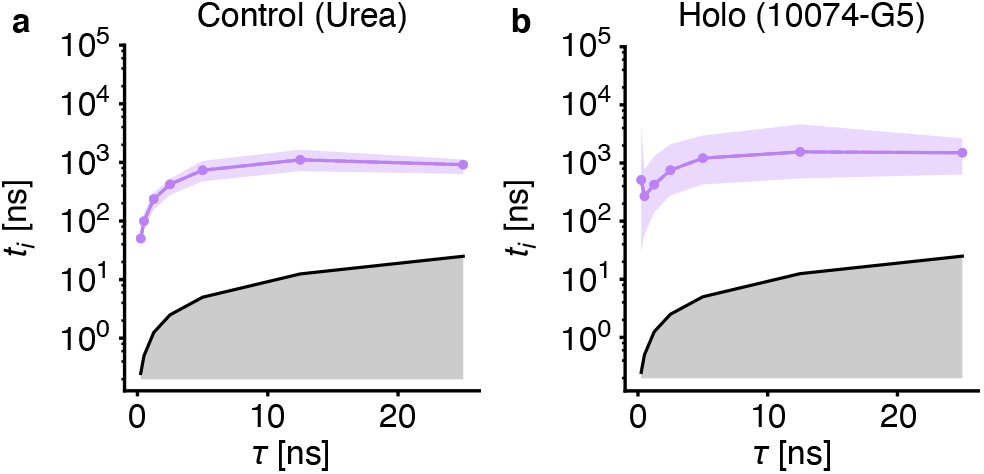
Relaxation timescale as a function of model lag time for **a** control (urea) and **b** holo (10074-G5) ensembles. Gray shaded areas indicate timescales the Koopman model can no longer resolve. Coloured shaded areas indicate 95th percentiles of the sample mean over all 20 models.

**Fig. A2.**
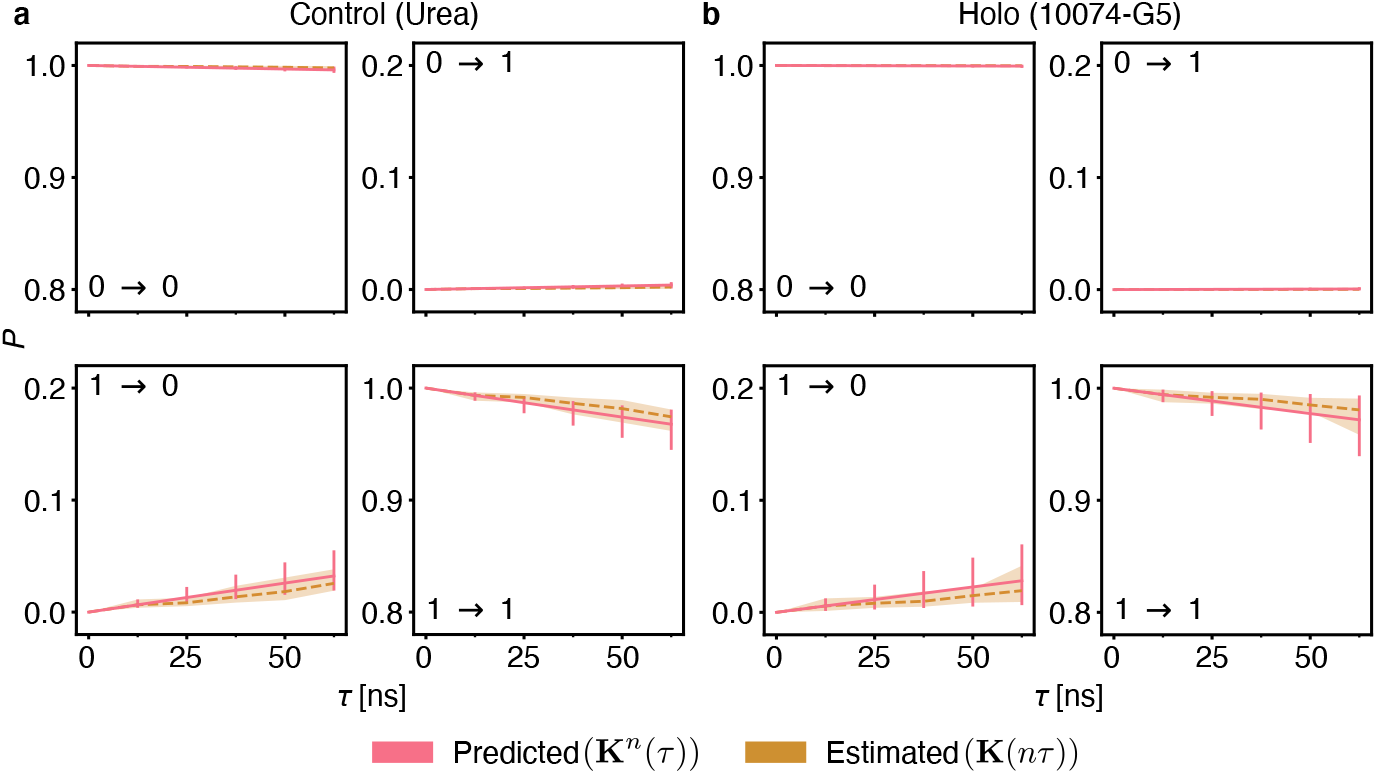
Chapman-Kolmogorov test for **a** control (urea) and **b** holo (10074-G5) ensembles. Each panel indicates the transition probability for one matrix entry for successive applications (predicted, red) and estimations (estimated, orange) of the Koopman matrix. Shaded areas and error bars indicate 95th percentiles of the mean over all 20 models.

**Fig. A3.**
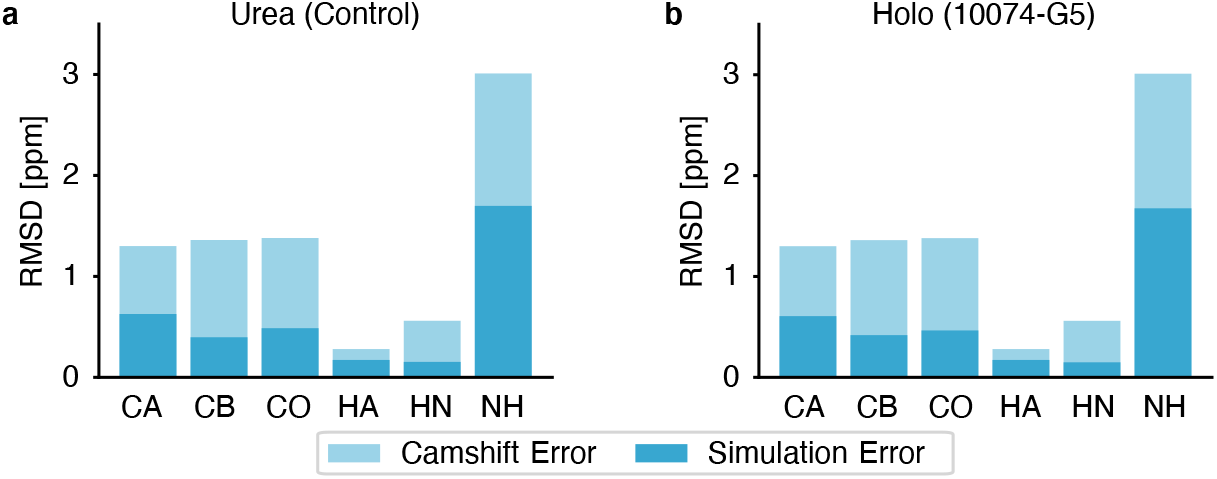
Root-mean-square deviations between experimentally determined NMR chemical shifts[5] and those back-calculated using CamShift[33] for **a** control (urea) and **b** holo (10074-G5) ensembles. Light shaded areas indicate the intrinsic error of the CamShift predictor.

**Fig. A4.**
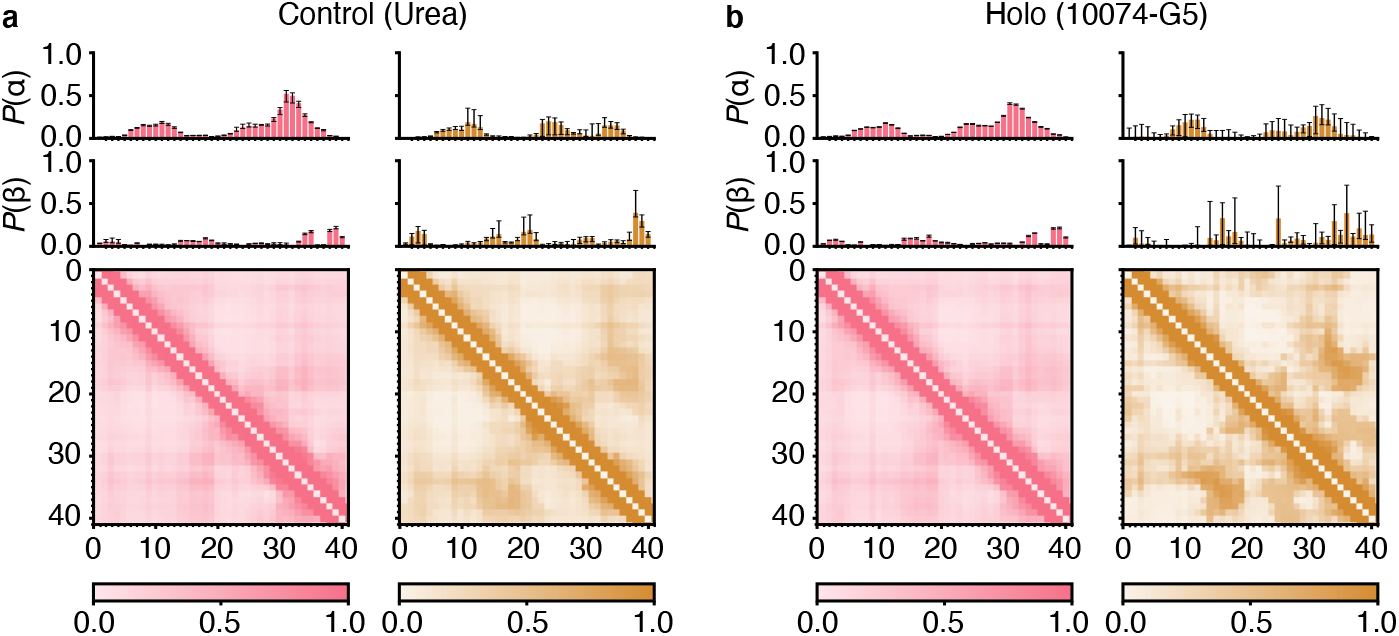
Structural properties of the **a** control (urea) and **b** holo (10074-G5) ensembles. Top panels indicate α-helical and β-sheet contents over all residues as calculated using DSSP[67]. Bottom panels show heavy-atom contact probability maps with a cut-off of 0.8 nm. Error bars indicate 95th percentiles of the bootstrap sample of the mean over all 20 models.

**Fig. A5.**
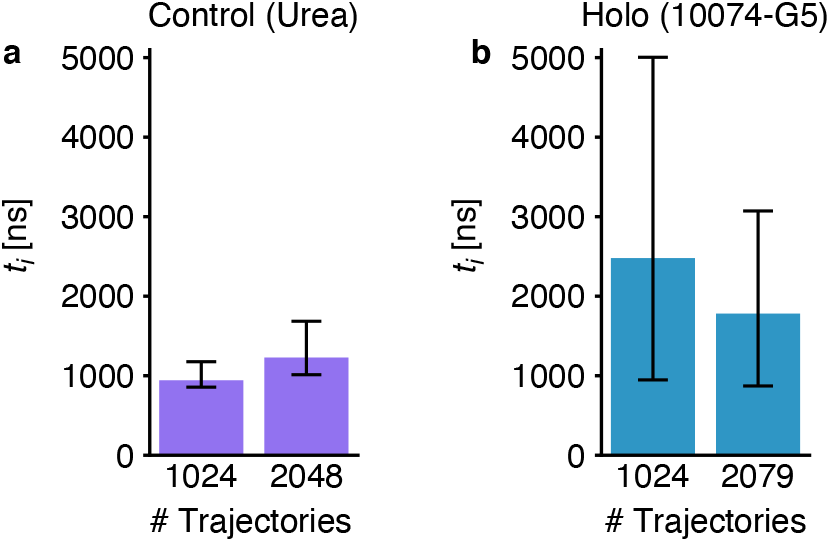
Dependence of the relaxation timescales on the number of trajectories used to build the model. Errorbars indicate 95th percentiles of the bootstrap sample of the mean over the first 5 models.

**Fig. A6.**
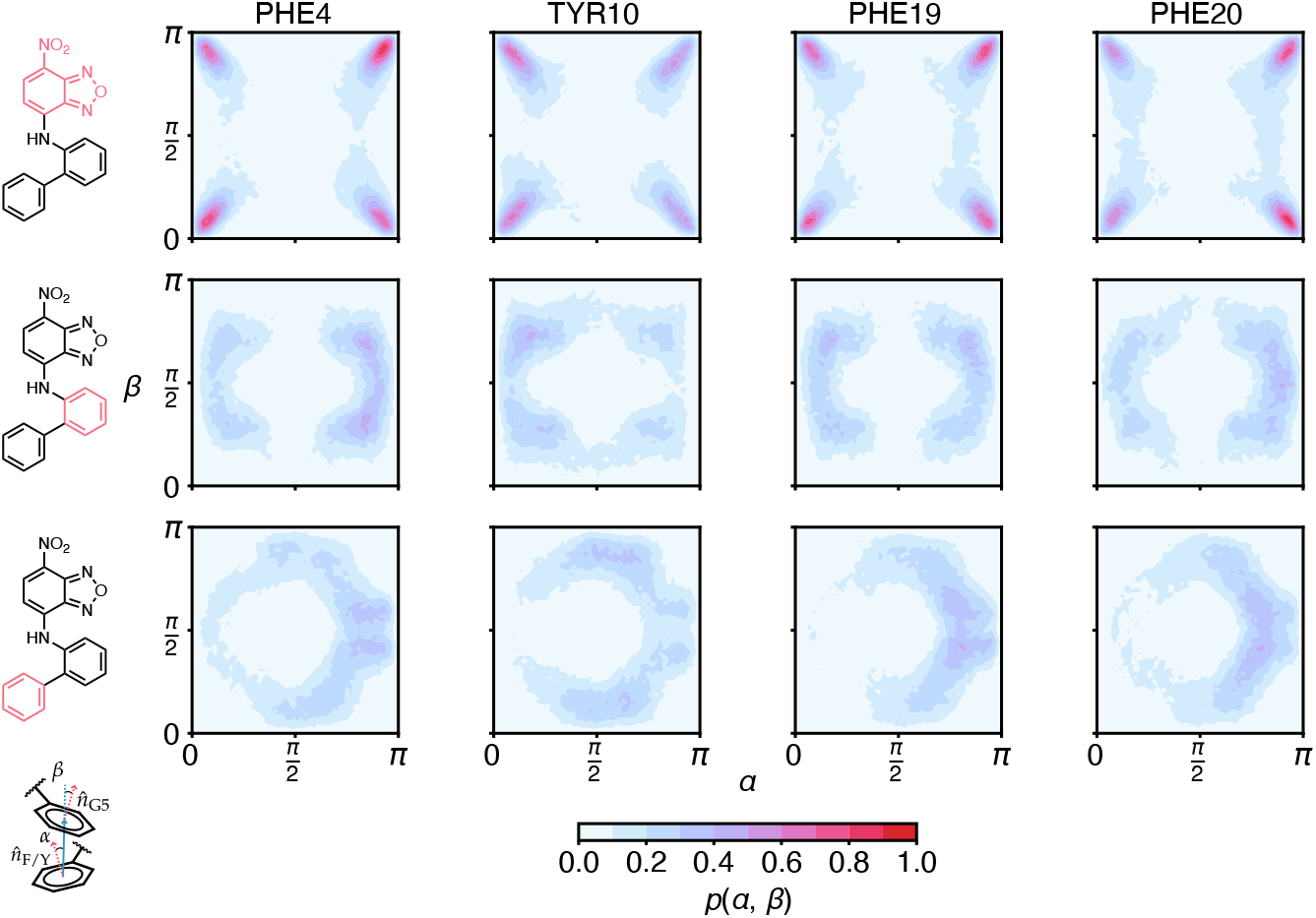
Anisotropy of π-π molecule — aromatic side chain stacking interactions for the holo (10074-G5) ensemble. *α* is the stacking angle between the inter-aromatic distance vector and the aromatic side chain normal vector, while *β* is the angle inter-aromatic distance vector and the normal vector of the small molecule aromatic system[35]. Distributions show the density under the condition that the distance between both groups is below 0.6 nm.

